# Competitive evolution of H1N1 and H3N2 influenza viruses in the United States: A mathematical modeling study

**DOI:** 10.1101/2021.09.30.462654

**Authors:** Chaiwat Wilasang, Pikkanet Suttirat, Sudarat Chadsuthi, Anuwat Wiratsudakul, Charin Modchang

## Abstract

Seasonal influenza causes vast public health and economic impact globally. The prevention and control of the annual epidemics remain a challenge due to the antigenic evolution of the viruses. Here, we presented a novel modeling framework based on changes in amino acid sequences and relevant epidemiological data to retrospectively investigate the competitive evolution and transmission of H1N1 and H3N2 influenza viruses in the United States during October 2002 and April 2019. To do so, we estimated the time-varying disease transmission rate from the reported influenza cases and the time-varying antigenic change rate of the viruses from the changes in amino acid sequences. By incorporating the time-varying antigenic change rate into the transmission models, we found that the models could capture the evolutionary transmission dynamics of influenza viruses in the United States. Our modeling results also showed that the antigenic change of the virus plays an essential role in seasonal influenza dynamics.

## 1. Introduction

The spread of the influenza virus is a major public health threat that leads to mortality, hospitalization, and an economic impact. Each year, there are one billion influenza cases globally, including 3 to 5 million cases of severe illness^1,2^ and at least 200,000 to 600,000 recorded deaths^3–6^. In the US, seasonal influenza accounts for more than tens of thousands of deaths during annual epidemics^7^. Moreover, the economic burden of seasonal influenza in the US has been estimated at more than ten billion dollars in healthcare and social costs^7^. The influenza epidemics recur annually and are very challenging to control. These recurrent epidemics are usually governed by a complex interaction of evolutionary and transmission dynamics of influenza viruses^1,6^. A better understanding of the evolutionary transmission dynamics of the virus could provide insights to mitigate the burden of seasonal influenza.

Influenza is caused by a rapidly mutating virus, which is an enveloped negative-strand RNA virus belonging to the family of *Orthomyxoviridae*^6,8,9^. The influenza viruses are classified into types A–D, where influenza type A is the most pathogenic^8^. Among influenza type A, different viral subtypes have been continuously reported. However, the subtype A(H3N2) and A(H1N1) are commonly found during epidemics each year^1^. In the US, these two influenza subtypes are reported almost every winter, usually from October to April. Seasonality is known to affect the transmission dynamics of the influenza virus^10–14^. A change in temperature and humidity throughout the year could profoundly affect the survival of these viruses. For example, in winter, the cold and dry weather can accelerate the growth of the virus and facilitate viral transmission^15^.

Vaccination is one of the most effective interventions to mitigate seasonal influenza epidemics^16,17^. Nonetheless, the annual vaccination may fail to fully protect vaccinated individuals due to antigenic changes of the viruses. Influenza viruses exhibit high genetic variability in which the viruses continually undergo an antigenic change through natural selection. For instance, the changes in the surface protein, called hemagglutinin (HA), help the viruses escape immune attacks^6,18,19^. The primary target of immunity against influenza is the HA proteins^7,20^. By changing the properties of antigenic sites in the HA proteins, the immune system may not recognize the virus and allows it to infect host cells^6,18,20,21^. Moreover, a high mutation rate has been observed in several studies due to the short generation times and large population sizes of the viruses ^22–24^. These evolutionary processes are believed to be a major driver of seasonal influenza transmission^20^.

In the US, influenza subtype A(H3N2) emerges and spreads more frequently than A(H1N1). Nevertheless, the two subtypes seem to be competitive for the susceptible hosts. Each year, the subtype virus that gets the highest fitness from mutation may spread predominantly in the population. In general, individuals infected with a single influenza strain will acquire immunity against that specific strain^16,22^. Partial or cross-immunity to other subtypes is also observed^15,25–29^. However, the level of cross-immunity is weakening with an increasing antigenic change of the influenza virus^2^. Therefore, co-circulation of the two influenza subtypes may result in cross-immunity.

Mathematical models, specifically the compartmental frameworks, have been widely used to investigate the transmission dynamics of influenza viruses. Most of the previous studies focused on the impact of seasonality on influenza transmission dynamics^12,13,30–33^. Subsequently, the evolution of these viruses has increasingly been studied as an important source of disease burden^6^. For instance, in 2014, J. B. Axelsen and co-workers developed a model to examine the role of climatic drivers and viral evolution on influenza dynamics^30^. The model also integrated an antigenic evolution process of the influenza virus by adding a new strain pulse to the year that a drastic antigenic change occurs. However, the model did not consider the evolutionary dynamics of the influenza virus. Later, in 2018, X. Du et al. developed advances in mathematical modeling to incorporate such antigenic changes. Their proposed model was designed specifically for seasonal H3N2 influenza to predict influenza incidence in the US^1^. Incorporating information on antigenic change into transmission models can accurately improve epidemiological forecasts. Although such models may fit the data accurately, their model does not consider the competition between different subtypes of the virus. Indeed, the co-circulation and co-transmission of influenza subtypes may result in different transmission characteristics^15,34^.

In this study, we investigated the impact of antigenic change on transmission dynamics of seasonal influenza. We, therefore, proposed epidemiological models for seasonal influenza A(H3N2) and A(H1N1) viruses that incorporate the antigenic changes of the viruses. The models integrated the data on the changes of amino acid sequences of HA proteins in epitope sites into the transmission models. To do so, we first estimated the time-varying antigenic change rate of each influenza subtype using the sequence data. We then incorporated this antigenic change rate into the transmission models to investigate the transmission dynamics of seasonal influenza in the US. We utilized long-term influenza surveillance data^35^ from October 2002 to April 2019 in the US to fit the models. Finally, we demonstrated that antigenic changes of the virus could play an important role in changing seasonal influenza transmission dynamics at the population level.

## 2. Materials and Models

### 2.1. Influenza surveillance data

The influenza surveillance data used in this study were obtained from the FluView of the US Centers for Disease Control and Prevention (CDC)^35^. The US CDC collects seasonal influenza activity data from both the US World Health Organization (WHO) Collaborating Laboratories and National Respiratory and Enteric Virus Surveillance System (NREVSS) laboratories. The US WHO and NREVSS laboratories report the total number of weekly respiratory specimens tested and the number of weekly influenza-like illness (ILI) cases that tested positive for influenza types A and B to the US CDC. Besides, the US WHO collaborating laboratories also report influenza subtypes A(H1N2) and A(H3N2). An ILI case was defined as a patient with a fever greater than 38 C° and a cough or sore throat^36,37^. Following X. Du et al.,^1^ the weekly influenza cases were computed as the product of the ILI incidence rate, the proportion of ILI samples that tested positive for influenza, the subtype proportion, and the US population size. The population data were downloaded from the US Census Bureau^38^. The monthly influenza cases from October 2002 to April 2019 were obtained by aggregating the weekly data.

### 2.2. Influenza amino acid sequences data

Amino acid sequences of influenza A(H3N2) and A(H1N1) were downloaded from the Global Initiative on Sharing Avian Influenza Data (GISAID)^39,40^. To analyze the HA protein sequences, whole sequences were aligned using Multiple Sequence Comparison by Log-Expectation (MUSCLE)^41,42^. The MUSCLE program uses FASTA files containing amino acid sequences as an input and provides the aligned amino acid sequences. Unknown amino acids were replaced by gaps. Aligned sequences were then edited using ClustalW, and the incomplete sequences were manually removed^43,44^. Since the major target of immunity against influenza is located in the epitope sites of the HA proteins^6,7,20^, we used only the amino acid sequences encoding the epitope sites of these proteins to analyze the antigenic change of the virus.

### 2.3. Evolution and transmission model with no competition

To investigate the evolution and transmission dynamics of influenza, we constructed two transmission models (for H3N2 and H1N1) that explicitly integrate the virus evolutionary dynamics (**Figure 1**). These models were used as baseline models to compare with competitive models. **Figure 1(A)** illustrates the evolutionary transmission model with no competition. In this model, the population was divided into three epidemiological classes, namely, susceptible, infectious, and recovered. The susceptible individuals (*S*) can be infected if they contact an infectious individual. After being infected, susceptible individuals move to the infectious class at rate *β_i_*(*t*)*I/N*, where *β_i_*(*t*) is a time-dependent transmission rate, and *N* is the total population size. The infectious individuals *(I)* recover from the disease after the infectious period (1/*γ*) and then are moved to the recovered class. Individuals in the recovered class are immune to the virus and cannot be infected. The model also integrates an effect of the virus antigenic change represented by an antigenic change rate (*ε_i_*(*t*)) as described in eq. (13). We assumed that recovered individuals (*R*) could become susceptible again at rate *ε_i_*(*t*)*R*. The evolutionary and transmission dynamics of influenza are described by the following ordinary differential equations (ODEs):

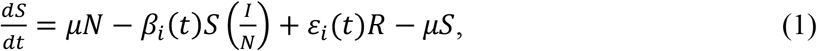

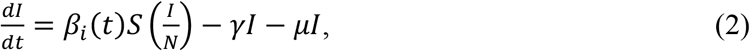

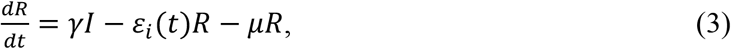

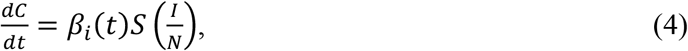

where *S, I, R,* and *C* represent the number of susceptible, infectious, recovered, and cumulative infected individuals, respectively. In this model, we assumed that the newborns are susceptible, and the number of natural births and deaths are balanced at a rate *μ* so that the total population size is constant. The parameters descriptions and the values used in the model are summarized in **Table 1**.

**Figure 1.**
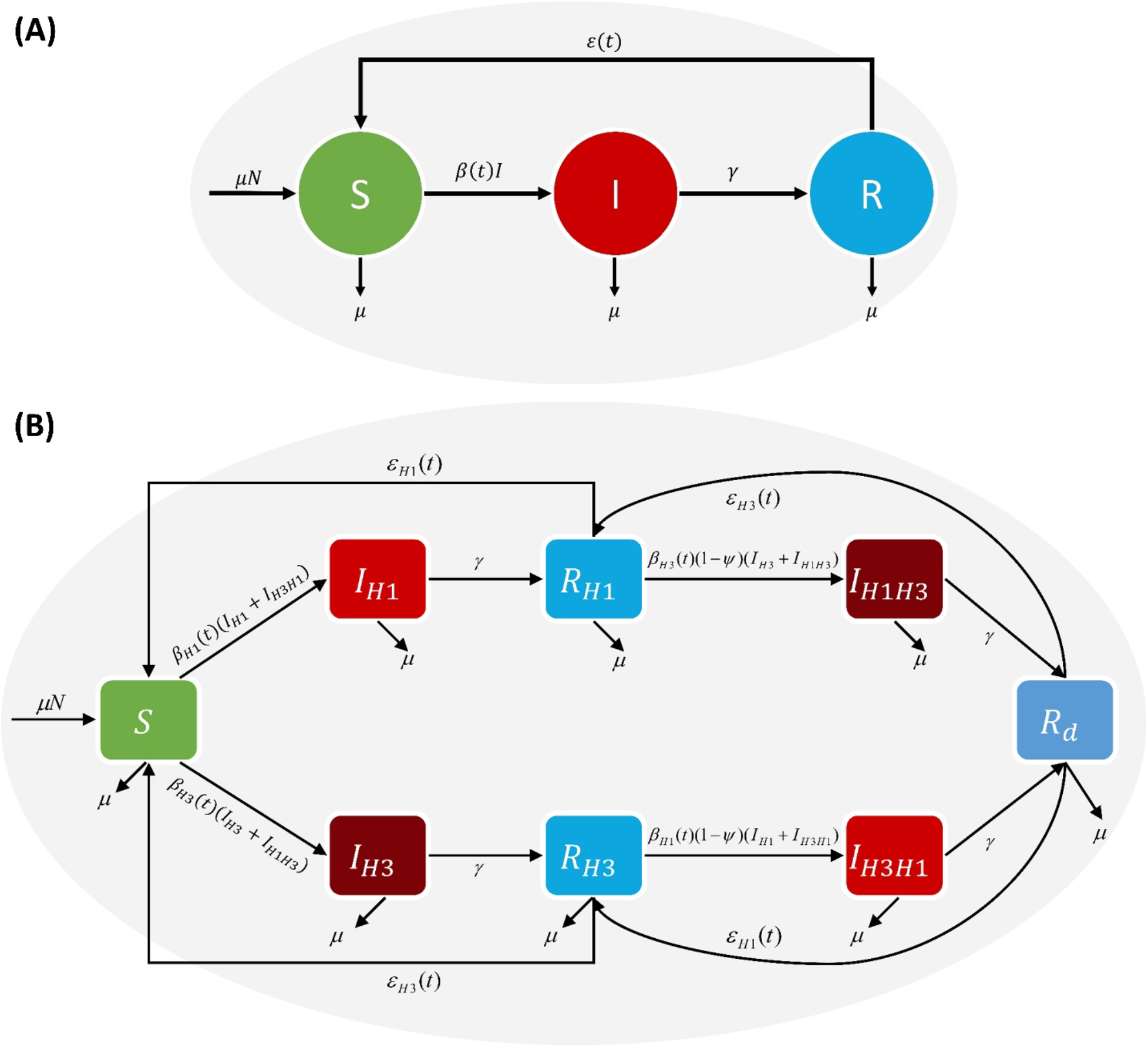
Schematic of the evolutionary transmission model. **(A)** The flow diagram of the evolutionary transmission model of the A(H1N1) and A(H3N2) influenza viruses with no competition and **(B)** with the selective competition.

**Table 1.** Model parameters and their values.

| Parameter | Definition | Value | References |
| --- | --- | --- | --- |
| $\beta_i(t)$ | Monthly transmission rate of influenza subtype $i$<br>= [H3N2 or H1N1] | [0.001-10] | Estimated |
| $\gamma$ | Recovery rate | 1/5 per day | 45-47 |
| $\mu$ | Natural birth and death rate | 1/67 per year | 3 |
| $\epsilon_0$ | Natural waning rate of immunity | 1/20 per year | Estimated |
| $\Psi$ | Cross-immunity induced by primary infection | 0.7 | 15,48,49 |
| $\alpha$ | The steepness of antigenic change | 2 | Assumed |
| $N$ | Population size | $3.25 \times 10^8$ | 38 |

### 2.4. Evolution and transmission model with selective competition

Several influenza virus strains can co-circulate in a population leading to selective competition in susceptible hosts^7,22,25,50,51^. The co-circulation of influenza viruses could result in cross-immunity and mediate the ecological competition between different strains. Consequently, for the modeling pursued here, we constructed an evolutionary transmission model considering cross-immunity and selective competition of the A(H3N2) and A(H1N1) influenza viruses. The schematic for the model with selective competition is shown in **Figure 1(B)**. Similar to the model with no competition, the population was categorized into three different groups, namely. However, each group of individuals here can be infected with either A(H1N1) or A(H3N2), indicated by the subscripts H1 and H3, respectively. Individuals recently recovered from the infection of one stain develop partial protection and hence are less susceptible to another strain. In our equations, we use the order of subscripts to indicate the order of infection. For example, *I_H3H1_* represents a group of individuals who had recently recovered from the infection of A(H3N2) but is currently infected with A(H1N1). *R_d_* represents a group of individuals that are immune to the two virus strains. The evolutionary transmission model with selective competition can be described by the following system of ODEs:

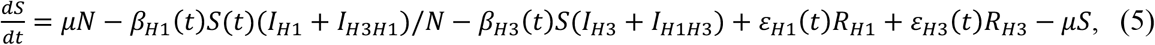

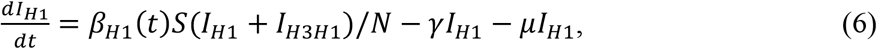

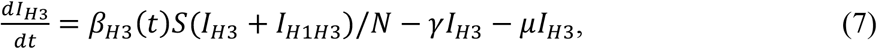

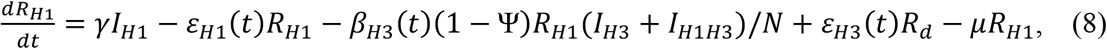

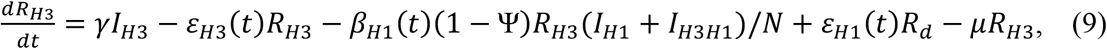

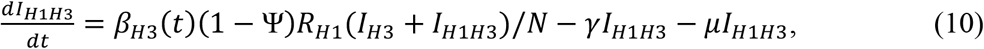

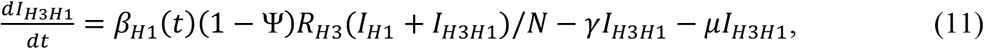

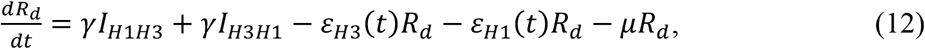

where *β*_*H*1_(*t*) and *β*_*H*3_(*t*) is the time-varying transmission rate of infectious individuals who are currently infected with A(H1N1) and A(H3N2), respectively. Ψ is the cross-immunity between the two influenza subtypes. *ε*_*H*1_ and *ε*_*H*3_ are the antigenic change rate of the A(H1N1) and A(H3N2) strain, respectively.

### 2.5. Estimating the time-varying antigenic change rate

Seasonal influenza viruses can better survive and spread in a specific environment, especially in winter^30,52^. The recurrence of seasonal influenza epidemics thus frequently occurs between October and March. To consider the effects of seasonality, the period from October 1 to March 31 was assigned as the high transmission season (HTS), while the remaining period was the low transmission season (LTS)^1^. We compared the amino acid sequences of both influenza subtypes between two consecutive seasons to estimate the antigenic change rate.

To investigate the evolutionary dynamics of the seasonal influenza virus, we used amino acid changes in the epitope as a measure of the antigenic distance between the two strains. Once the virus infects the host cells, the antibody attacks the antigen by binding to HA proteins located on the viral surface^18^. This part of an antigen memorized by the immune system is called epitope^3,53^. The mutation within epitope regions could, thus, weaken the immune system^19^. Consequently, the virus can enter and infect the host cell. Epitope regions are categorized differently according to influenza virus subtypes^6,54–57^. The amino acid residues of epitope regions are summarized in **Table S1** and **Table S2** in the Supplementary Information.

To estimate the antigenic change among influenza viruses, we considered aligned amino acid sequences as character strings and computed the number of amino acid differences within all five epitopes, known as the Hamming distance^58^. The Hamming distances were calculated by comparing the viral sequences in the current and previous seasons. However, as the numbers of viral sequences recorded in each season are different, we first sampled 10^6^ random pairs of sequences between seasons to avoid sampling biases. Subsequently, we calculated the Hamming distance for each pair of amino acid sequences. We then computed *P_all-epitopes_*, which is a fraction of the number of amino acid differences within all five epitopes and the total number of amino acids presented^59^.

Our model incorporated the antigenic change rate into the transmission model to investigate how antigenic change affects transmission dynamics. We used the functional form for the time-varying antigenic change rate as shown in eq. (13), which allows us to explore a range of antigenic change rate regimes: when *P_all-epitope_* = 1, the antigenic change rate is very high (as might be the case if antigenic change between two influenza virus strains was completely changed), and when *P_all-epitope_* = 0, the antigenic change rate is low. The time-varying antigenic change rate, *ε*(*t*), is given by:

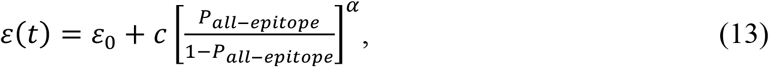

where *ε*_0_ represents the naturally waning rate of immunity. Parameters *α* and *c* were estimated such that the resulting cumulative cases derived from the simulations match the influenza surveillance data. We varied the antigenic change rate and simulated the H1N1 influenza model before the pandemic season of 2009–2010. The values of *ε*(*t*) that the model can search for are uniformly distributed in the interval [0, 1] with 0.001 resolution. We recorded only the values of the antigenic change rate that yield the best simulation results. Subsequently, we calculated the constant parameter *c* according to eq. (13). Simultaneously, we varied *α* the value of steepness changes. The estimated parameters are *α* = 2 and *c* = 5.9911×10^-2^ as they minimize the difference between simulated and reported data.

### 2.6. Estimating the time-varying transmission rate

In this study, we employed the parameter estimation method developed by P. Suparit et al.^60^ to estimate the time series of monthly transmission rates. The monthly influenza cases in the US starting from October 2002 to April 2019 were used to estimate the time series of monthly transmission rates. Briefly, we divided the model simulation into several consecutive monthly time window simulations. We varied *β*(*t*) ranging from 0.001 to 10 with the resolution of 0.01 and searched for the transmission rate that provides the best fit between the number of monthly simulated cases (*C_sim_*) and the number of monthly reported cases (*C_rep_*). The values of the monthly transmission rate that minimizes |*C_sim_* – *C_rep_*| were recorded in the time series of monthly transmission rates. We also recorded the number of individuals in each epidemiological class for the current monthly time window simulation. Subsequently, we used the number of individuals in the previous monthly time window as an initial condition for the next monthly time window simulation. The process was repeated until the complete time series of the monthly transmission rates was obtained.

### 2.7. Simulation details

To generate the long-term seasonal influenza dynamics, the infection was initiated by randomly selected seed infections and run for a period of 100 years to allow the system to reach equilibrium. We then used the averaged population in each compartment as the initial population of the models. For the simulation process, we used the Euler method to solve the sets of ODEs implemented in MATLAB R2019b.

## 3 Results

### 3.1. Seasonal patterns of influenza epidemics in the US

The monthly influenza cases from October 2002 to April 2019 (~17 years) reported in the United States are shown in **Figure 2** (see also **Figure S1**). The data shows the incidence of monthly influenza of two subtypes: subtype A(H3N2) and subtype A(H1N1). The outbreaks of influenza A (H3N2) and A (H1N1) usually appear between October and March. The numbers of cases are usually lower in the summer season (April to September). We, therefore, classified the period from 1^st^ October to 31^st^ March as a high transmission season (HTS) and the period from 1^st^ April to 30^th^ September as a low transmission season (LTS). Thus, there were 17 HTSs and 16 LTSs in the time series. As illustrated in **Figure 2**, both subtypes of influenza viruses have co-circulated throughout the study period. Aside from the apparent annual cycles, epidemics of A(H3N2) and A(H1N1) usually occur at different magnitudes, especially during the year 2009 when the novel 2009 A(H1N1) influenza emerged ^47,61,62^. Furthermore, we observed a competitive pattern between influenza A(H3N2) and A(H1N1). The observed incidence data suggested that when influenza A(H3N2) prominently influenced the epidemic, influenza A(H1N1) would subside, and vice versa.

**Figure 2.**
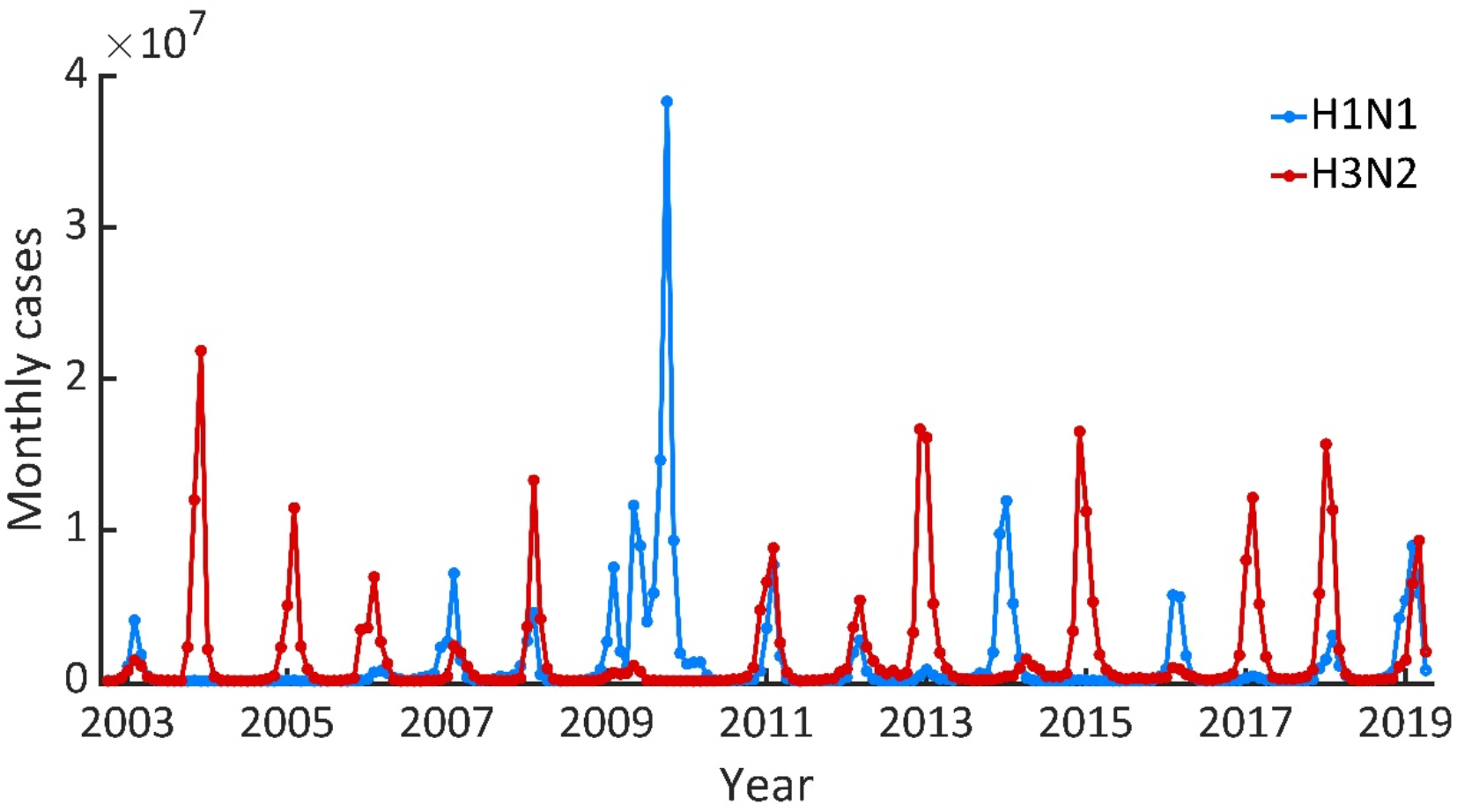
Influenza epidemics in the US. The monthly cases of influenza A(H1N1) and A(H3N2) in the US between October 2002 and April 2019.

### 3.2. Antigenic change rates of influenza virus

The recurrent influenza epidemics are a combined outcome of viral antigenic changes, interactions among co-circulating influenza viruses, transmission, and host immune recognition against the virus^6^. When influenza viruses undergo evolution, the immune system may not be able to recognize the viruses, allowing the viruses to infect host cells. Thus, previously infected individuals may eventually become susceptible to the viruses again^19^. To quantify the antigenic change rate, we calculated *P_all-epitopes_*, which is a measure of the antigenic change between two influenza virus strains^4,59^ (see Material and Methods for more details). The average *P_all-epitopes_* for both influenza A(H3N2) and A(H1N1) are shown in **Figure 3 (A)** and **(B)**, respectively. *P_all-epitopes_* were also compared with the monthly influenza cases. According to the results shown in **Figure 3(A)** and **(B)**, we found that the large epidemic curves usually occur after a significant antigenic change of influenza viruses. Specifically, the antigenic change of influenza A(H1N1) was around 80% in the 2008-2009 HTS, which resulted in the subsequent surge of the 2009 influenza pandemic (**Figure 3(B)**). This result suggested that *P_all-epitopes_* might be a potential early warning for a pandemic. We also found that the average *P_all-epitopes_* of A(H3N2) is less fluctuate than that of A(H1N1). The corresponding antigenic change rates of influenza A(H1N1) and A(H3N2) are illustrated in **Figure 3(C)** and **(D)**.

**Figure 3.**
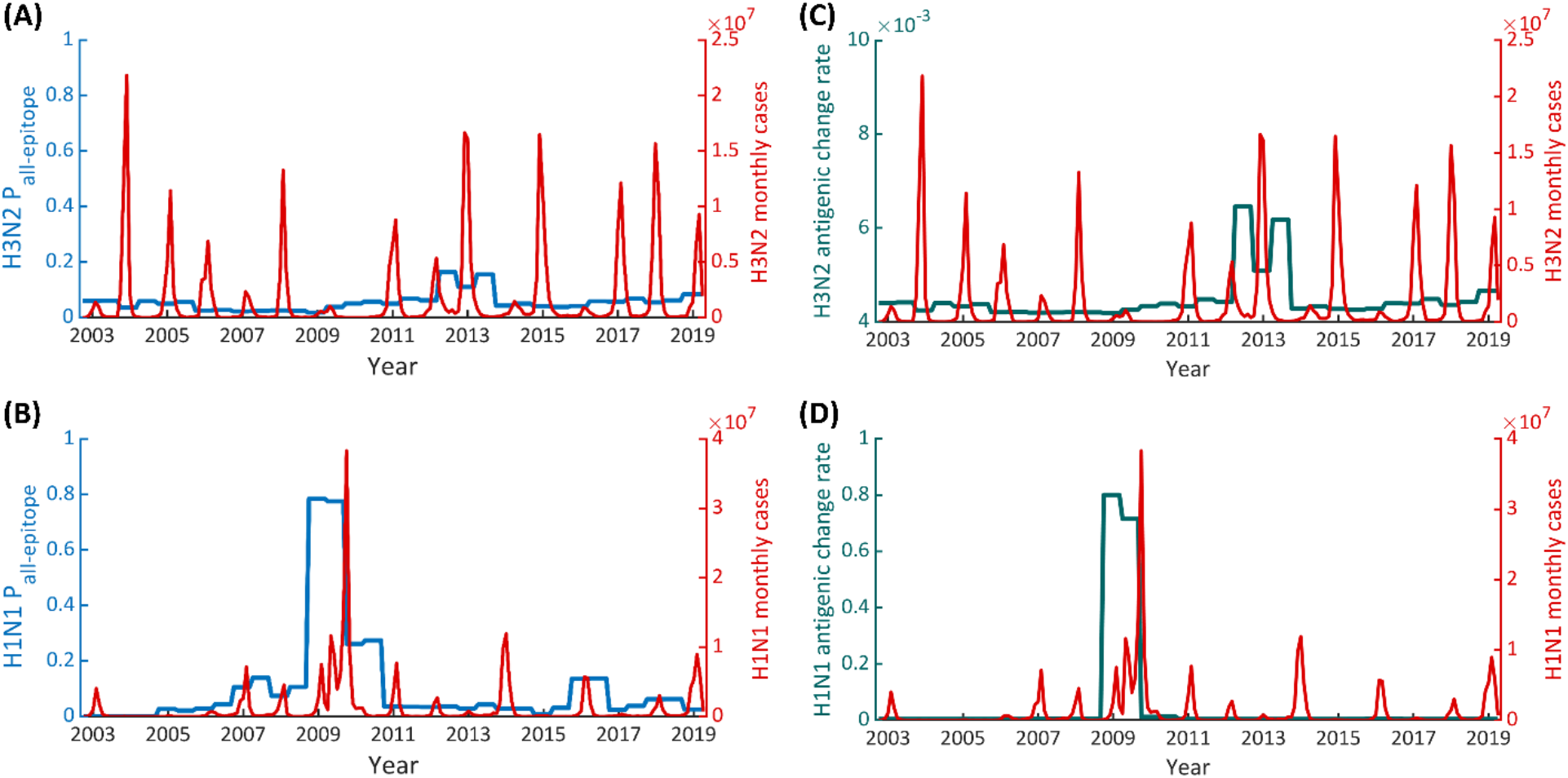
Average monthly *P_all-epitopes_* and antigenic change rates. *P_all-epitopes_* of influenza **(A)** A(H3N2) virus, and **(B)** influenza A(H1N1) virus, **(C)** the antigenic change rate of influenza A(H3N2) virus, and **(D)** influenza A(H1N1) virus.

### 3.3. Effect of the antigenic change of the influenza virus

We found that the estimated transmission rates from the two models are different. The estimated monthly transmission rates that provide the best fit of the two models are shown in **Figure 4**. The results illustrate that the transmission rates estimated from the model without the evolutionary dynamics increased after the major changes of influenza viruses in 2009. In contrast, there is no change in the post-2009 trend of the transmission rates estimated from the model incorporating the evolutionary dynamics. The transmission rate fluctuated between 0.11 and 0.31 per month during the years 2009 and 2019. There is also an approximately 2-fold increase in the oscillation amplitude of transmission rates when excluding the evolutionary dynamics in the model. These suggested that the antigenic change of the virus could affect their transmission dynamics. This effect was also observed in the transmission dynamics of the influenza A(H3N2) virus (**Figure S3** in the supplementary results).

**Figure 4.**
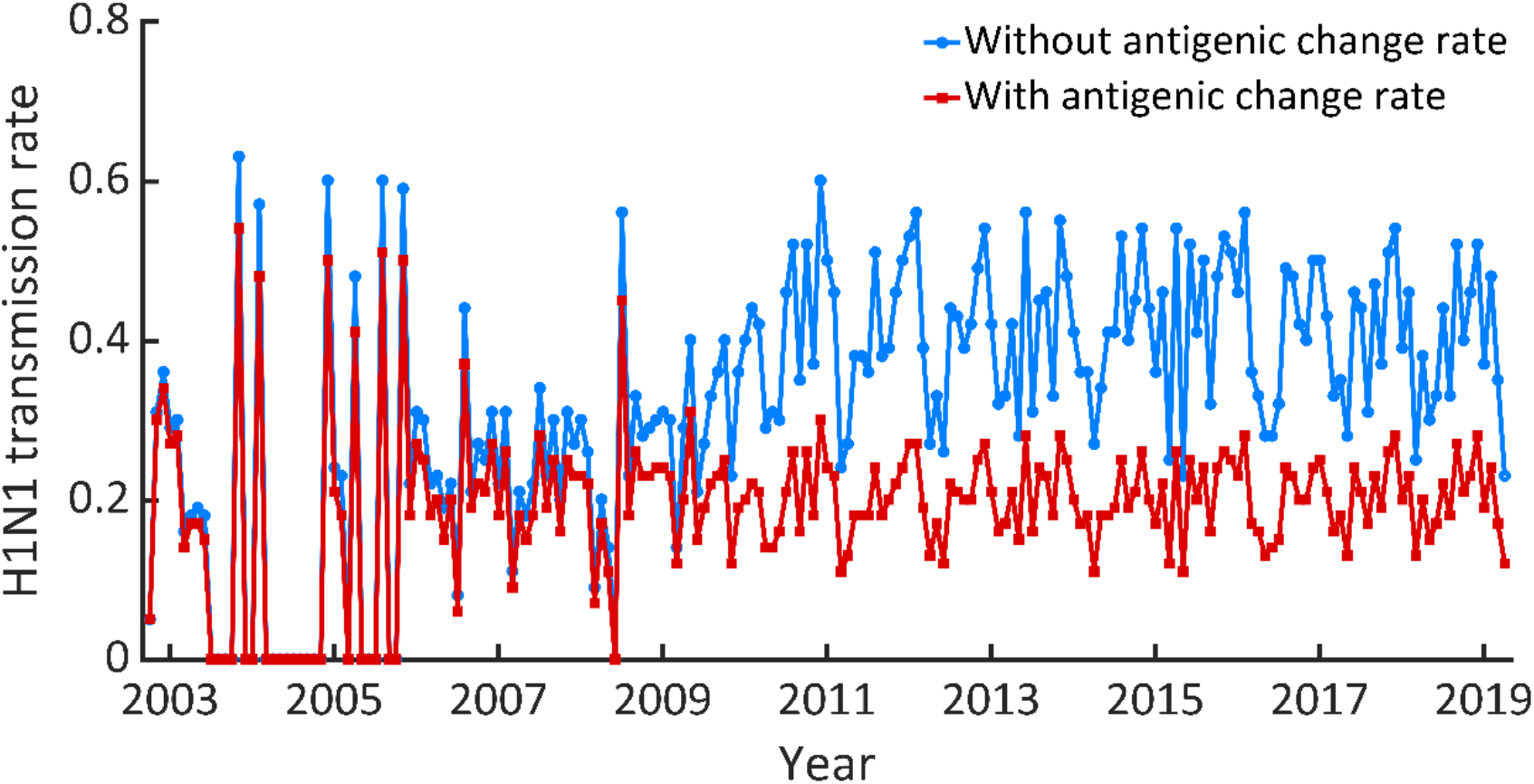
Comparison of H1N1 monthly transmission rates. The lines show the time series of monthly transmission rates of the A(H1N1) influenza virus during October 2002 and April 2019. The blue line shows the monthly transmission rate estimated using the model without antigenic change. The red line denotes the monthly transmission rate estimated by the model with the antigenic change of the virus.

### 3.4. Competitive evolution and transmission dynamics of the A(H3N2) and A(H1N1) influenza viruses

The infection of a certain influenza subtype in humans results in partial immunity to other subtypes of influenza viruses^22,63^. The empirical evidence suggests that the co-circulation of influenza viruses not only results in cross-immunity but also appears to mediate the ecological competition between the subtypes^7,22,25,50,51^. A competitive model was then constructed to investigate the combined co-evolution and co-transmission dynamics of the A(H3N2) and A(H1N1) influenza viruses. We also incorporated the cross-immunity between the two subtypes into our competitive model. The results illustrated that the models could faithfully reconstruct the influenza epidemics (**Figure S4** and **Figure S5** in the supplementary results).

We then further investigated the effects of cross-immunity between the two subtypes. The strength of cross-immunity between the two influenza subtypes is represented by the parameter Ψ. If Ψ = 0, the influenza subtypes are completely different antigenically. In contrast, if Ψ = 1, the two subtypes are antigenically indistinguishable^25^. **Figure 5** illustrates the effect of the cross-immunity on the transmission rates estimated by the three models. The transmission rate estimated from the competitive model with the cross-immunity Ψ = 0.7 was found in the range of 0.01–1.60 per month with an average value of 0.59. In contrast, the transmission rates estimated from the competitive model without cross-immunity (Ψ = 0.0) fluctuates in the range of 0.01-0.60 with an average value of 0.20 per month, which is close to the transmission rates estimated from the single strain model (H1N1 model).

**Figure 5.**
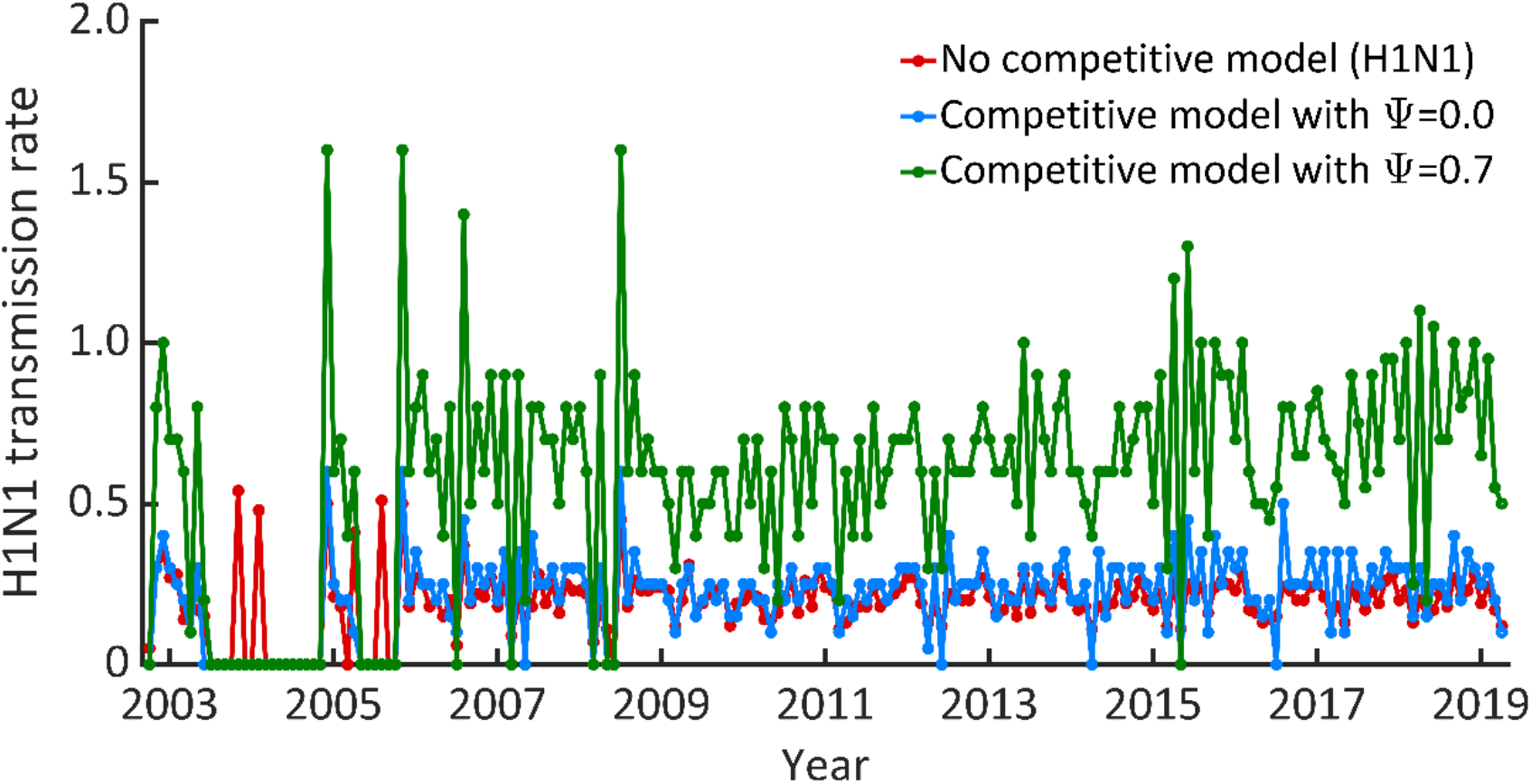
The time series of transmission rate estimated from the three different models. The red line shows the monthly transmission rate of the H1N1 influenza virus estimated from the model without antigenic change of the virus. The blue and green lines display the monthly transmission rate of the H1N1 influenza virus estimated from the competitive model without cross-immunity (Ψ = 0.0) and with cross-immunity Ψ = 0.7, respectively.

Comparing the proportion of immune individuals predicted by each model, we found the fraction of the immune population obtained from the non-competitive model is lower than that of the competitive one (**Figure 6** and **Figure S6(B)** in the supplementary results). Noticeably, at the end of 2009, the immunity levels of influenza A(H1N1) from the three modeling scenarios were predicted at 7.94%, 12.82%, and 20.10%, respectively. Regarding the seroprevalence of influenza-specific antibodies, we found that the competitive model with cross-immunity provides the proportion of the immune population closest to the observed data at 21.0%^64,65^.

**Figure 6.**
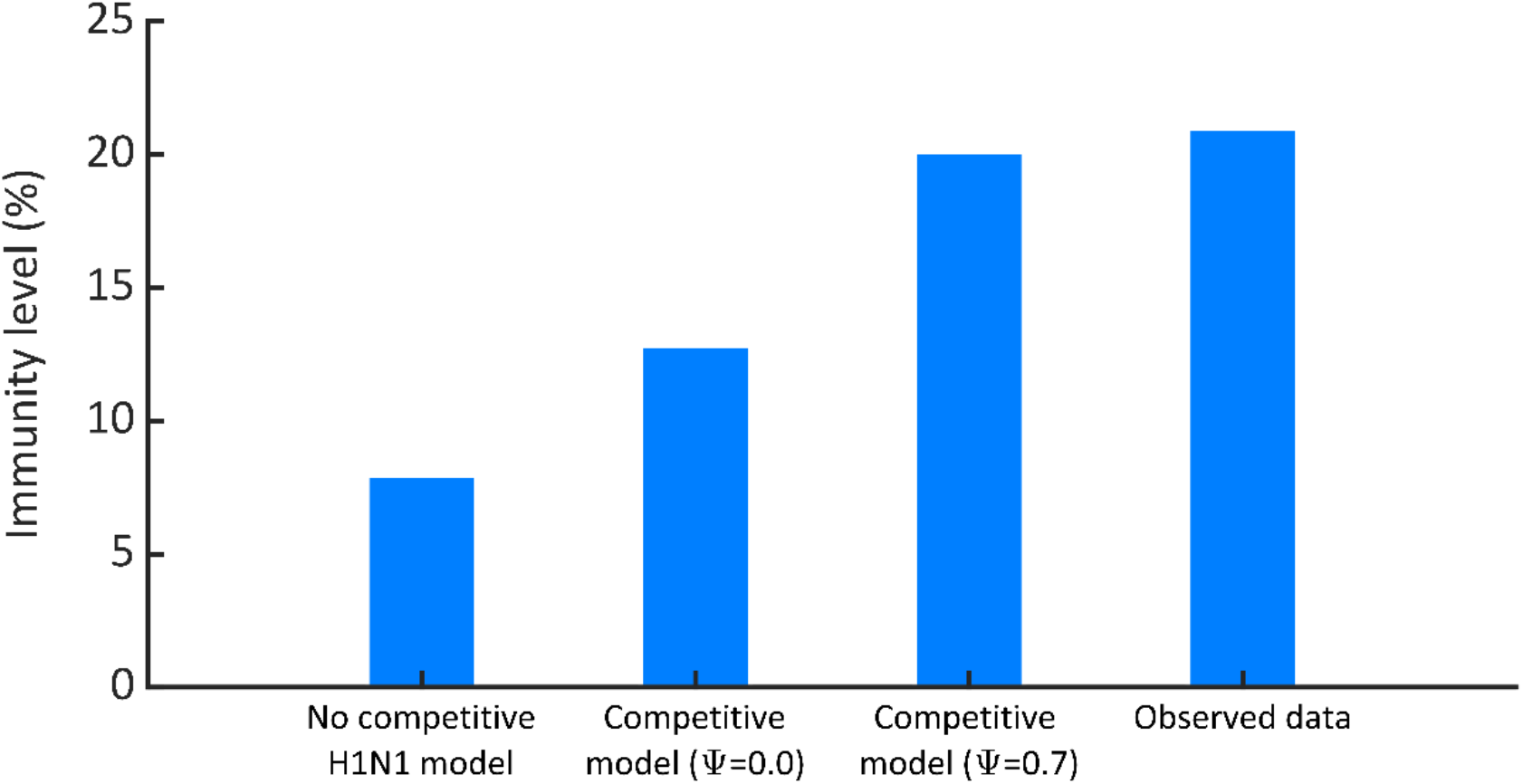
Immunity level of influenza A(H1N1). The immunity levels of influenza A(H1N1) estimated from the H1N1 model, the competitive model without cross-immunity, and the competitive model with Ψ =0.7 are 7.94%, 12.82%, and 20.10%, respectively. The seroprevalence of influenza observed by S. M. Zimmer et al.^64^ was approximately 21.0%.

## 4. Discussion

Influenza epidemics have been observed annually, resulting in significant morbidity and mortality. The recurrence of the influenza epidemic stems from several possibilities^6^. Seasonal factors are known to affect the transmission dynamics of the influenza virus^10–14^. However, one of the most significant factors that drive the annual influenza epidemic is the evolution of the virus^1,19,58^. Thus, the integration of the antigenic change of the virus into the epidemic model is necessary. To capture the transmission dynamics of the seasonal influenza virus in the US, we proposed mathematical models incorporating the antigenic change of the virus. The models were employed to reconstruct the long-term influenza epidemics in the US from October 2002 to April 2019. We also compared competitive evolutionary transmission models of influenza A(H1N1) and A(H3N2) viruses with and without cross-immunity.

To investigate the interplay between the epidemic dynamics and the evolutionary dynamics of the influenza virus, we compared the virus antigenic change rate with the monthly influenza incidence (**Figure 3**). The antigenic change rate of influenza A(H3N2) increased a little during the season 2013 (October 2012 to April 2013), which might contribute to the large outbreak in the US in that season, resulting in approximately 30 million influenza illnesses^66^. For influenza A(H1N1), the antigenic change rate drastically increased during October 2008 and April 2009, consequently causing the 2009 A(H1N1) influenza pandemic^67^. Over this period, we observed that the antigenic change rate of influenza A(H1N1) considerably increased from 0.01 to 0.80. Thus, the emerging strain could replace the existing stains rapidly^68^.

Several studies have examined the impacts of seasonal factors (e.g., temperature and humidity) on the transmission of seasonal influenza^11–14,25^. Few studies have attempted to bridge the gap between the antigenic change of the virus and the transmission model^1^. Previously, this was modeled by adding a constant rate to the year that a drastic antigenic change occurs to represent the effect of the influenza virus evolution^30^. However, in this work, we intended to link antigenic changes of the influenza virus to disease transmission models. Specifically, we investigated the effect of antigenic change on seasonal influenza epidemics by incorporating the antigenic change rate into the transmission model. We also validated our models by comparing the modeling results with the reported data. We found that our models could reconstruct the observed epidemic dynamics for both high transmission and low transmission seasons for the two subtypes.

Furthermore, the antigenic change rate included in the models can describe the transmission dynamics of seasonal influenza. To interpret these findings, we compared the monthly transmission rates estimated from the models with and without antigenic change of the influenza virus. We observed significant differences in the transmission rates obtained from these two models. The transmission rate estimated from the model incorporating antigenic change rate fluctuated around a certain constant level even though there was a drastic change in the genetic sequence of the influenza virus in 2009. In contrast, for the model without the antigenic change rate, the fluctuation of the estimated transmission rate was shifted up after the year of the pandemic. The increase in transmission rate may result from the large outbreaks in 2009, which drastically depleted the susceptible pool. Consequently, the model compensates by increasing the transmission rate to generate the same number of reported cases. This means that either human behavior or viral transmission characteristics would have abruptly changed. However, there is no evidence that those events occurred, unlike the current COVID-19 pandemic^69,70^. Since the transmission rate depends on human behavior, it is unlikely to change after the 2009 pandemic. These results highlight the importance of incorporating the antigenic change rate into the model. The models incorporating evolutionary information of seasonal influenza virus could explain in more logical prediction on virus characteristics such as transmissibility and immunity. Thus, to accurately reconstruct the dynamics of seasonal influenza transmission, the antigenic change of the virus should be considered in transmission models.

Indeed, people can be infected by multiple strains of the influenza virus. We, therefore, constructed competitive transmission models to investigate the combined co-evolution and co-transmission dynamics of the A(H3N2) and A(H1N1) influenza viruses. Our findings indicated that population susceptibility is associated with the antigenic change of the virus and the strength of cross-immunity. The antigenic change of the virus replenishes the pool of susceptible individuals, which increases population susceptibility, whereas cross-immunity can reduce population susceptibility^71^. The antigenic variation of the influenza virus may allow the virus to escape the existing protective immunity and cause new epidemics. This antigenic change can accommodate the system dynamics, e.g., increased population susceptibility, and allows abrupt increases in population susceptibility in response to epidemic surges. In addition, we also found that there was a competition between influenza subtypes via cross-immunity. Different values of estimated transmission in **Figure 5** illustrate the effect of cross-immunity. These indicate that cross-immunity could modulate the ecological interactions between co-circulating influenza viruses by increasing the transmission rate of the virus. If the cross-immunity level is high, the competition between the two subtypes becomes extensively strict^25^. This finding agrees with the conclusions from other studies^2,25,50,51^, which found that cross-immunity among influenza viruses would naturally lead to competition between subtypes. Besides, we also validated our competitive transmission models by comparing the level of immunity from model predictions with seroprevalence data. The competitive transmission model with cross-immunity predicted the immunity level at the end of 2009 to be 20.10%, which is close to the values found in other studies^64,65,72–74^.

Our models, however, made several simplifying assumptions: firstly, we assumed that the population is homogeneously mixed and that all individuals have an equal probability of transmitting the disease. Second, we ignore heterogeneity in disease transmissions such as sex, age structure, and geographical region. The individual heterogeneity in infectiousness, e.g., superspreading events, is known to affect the dynamics of disease transmission^75,76^ and stochastic extinction^76^. Finally, we did not consider human mobility in our model; incorporating these data into the model might improve the prediction power of the model^77,78^.

## 5 Conclusion

In summary, we constructed evolution and transmission models to investigate seasonal influenza transmission in the US. The models integrate the changes in amino acid sequences of HA proteins in epitope sites and the time-varying disease transmission rates. We found that the models with the information on antigenic change outperform the models without it. We also constructed the competitive models to investigate the competitive evolutionary transmission dynamics of the A(H3N2) and A(H1N1) influenza viruses. Our results suggested that cross-immunity could modulate the ecological interactions between co-circulating and naturally lead to competition between subtypes. Ultimately, investigating evolutionary change is an important aspect of developing a current understanding of the circulation and prediction mechanisms of influenza viruses.

## Supporting information

supplementary results

## Supporting information

S1 Text. Supplementary Information

## Acknowledgments

Chaiwat Wilasang is supported by the Science Achievement Scholarship of Thailand (SAST).

## Authors’ Contributions

CW participated in its design, performed the simulations, performed the analysis, and wrote the first draft. PS participated in analyze the results and edited the manuscript. SC and AW participated in its design, analyzed the results, and wrote the manuscript. CM conceptualized, participated in its design, analyzed the results, and wrote the manuscript. All authors read and approved the final manuscript.

## Availability of Data and Materials

The data supporting the findings can be found in the main paper.

