## supplementary results for "Competitive evolution of H1N1 and H3N2 influenza viruses in the United States: A mathematical modeling study"

**Figure S1** shows the seasonal pattern of influenza A (H3N2) and A (H1N1). The monthly cases are plotted against the seasons to observe the effect of seasonal factors. As can be seen, the outbreaks of both influenza subtypes usually appear between October and March each year, which is defined as a high transmission season (HTS). The numbers of cases are frequently lower in the low transmission seasons (LTS). However, in 2009, the 2009 A(H1N1) pandemic caused outbreaks with atypical seasonality and spread throughout the US.

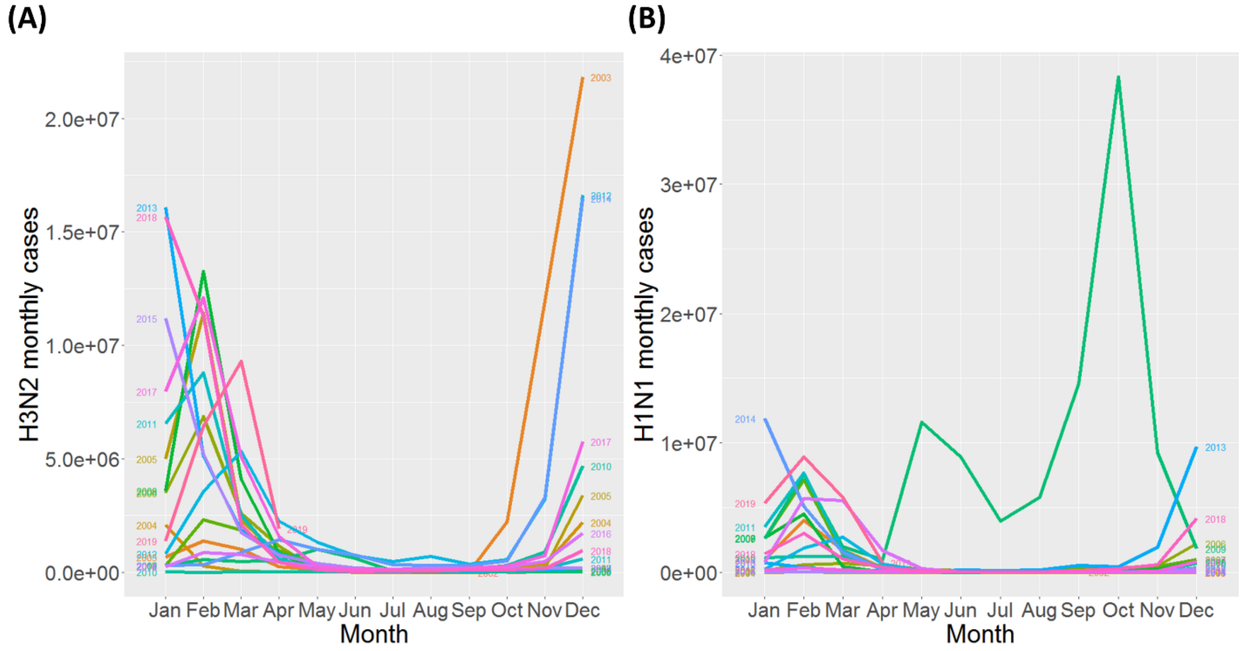


**Figure S1. Seasonality in the time series of influenza. (A)** A seasonal plot of influenza A(H3N2) monthly cases. **(B)** Seasonal plot of influenza A(H1N1) monthly cases. Lines show the yearly time series of incidence.

### Seasonal influenza transmission dynamics in the US

We found that the trends of monthly influenza cases generated from the model agree well with the surveillance data (**Figure S2**). During the pandemic season of 2008-2009, for example, the total number of influenza cases was predicted at 39.03 million, which is close to the number of reported cases at 38.3 million. We also employed the model to reconstruct the transmission dynamics of influenza A(H3N2) during the same period in the US. We found that the H3N2 model also provided a good fit to the observed data with R-square = 0.9998 (**Figure S2**). **Figure S2(A)** and **S2(B)** show the comparison between the modeling results and the surveillance data. We found that the trends of monthly cases generated by the influenza A(H3N2) and influenza A(H1N1) models agree well with the reported data with R-square = 0.9998 and 0.9989, respectively. The results suggested that the model incorporating the evolutionary rate could also capture the influenza epidemic data well. Our models can recreate the observed epidemic dynamics during both epidemic and non-epidemic periods for all subtypes.

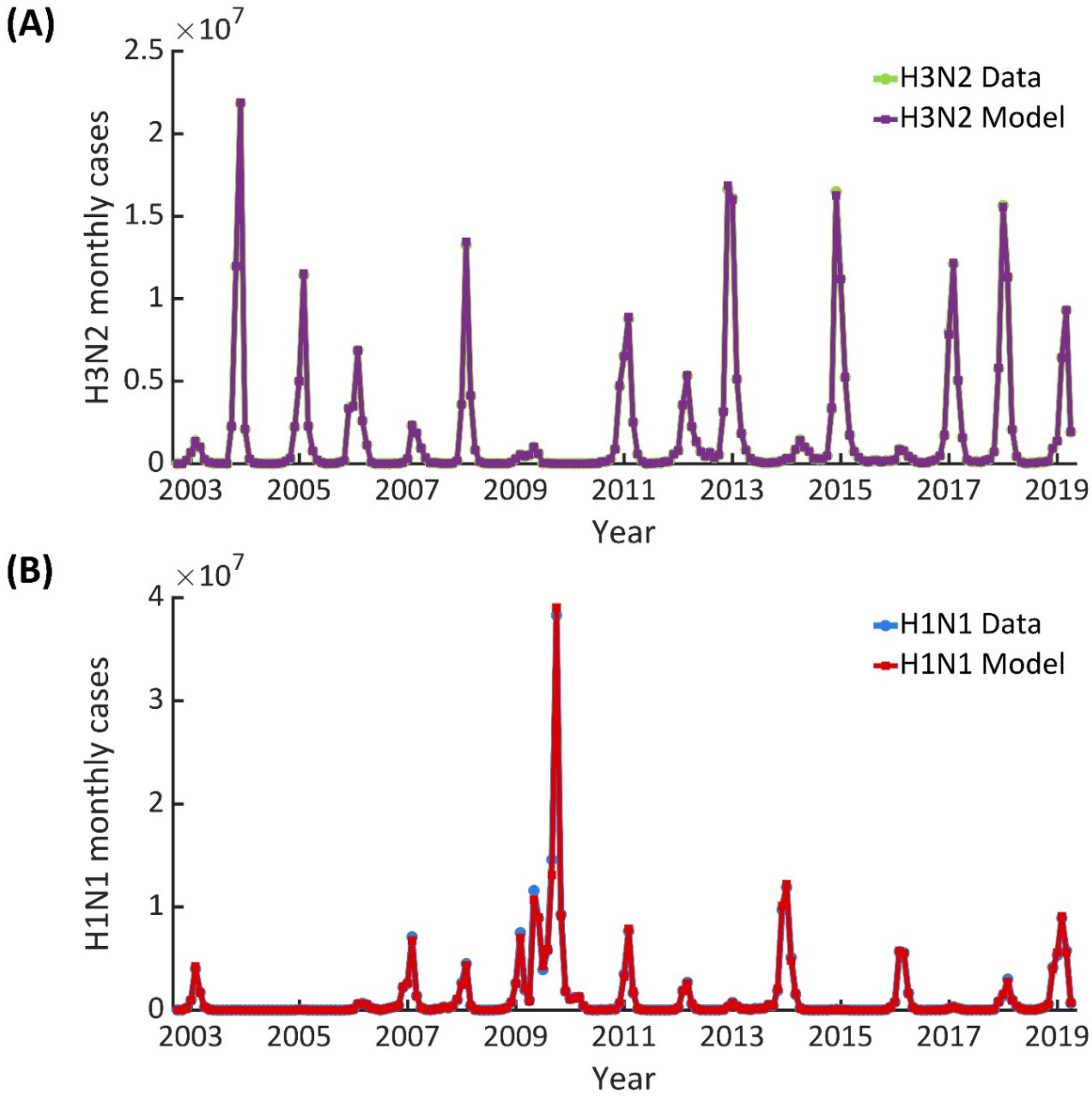

**Figure S2. Model fitting**. **(A)** Influenza A(H3N2) model. The model fit of the 16 years of historical influenza data (October 2002 to April 2019) correlates with the observed data with R-square = 0.9998. **(B)** Influenza A(H1N1) model. The model fit of the 16 years of historical influenza data (October 2002 to April 2019) correlates with the observed data with R-square = 0.9989.

To investigate the role of the evolutionary change of the influenza virus in the dynamics of influenza epidemics, we compared the estimated transmission rate of influenza viruses from two different models. **Figure S3** shows the estimated monthly transmission rates of influenza A (H3N2). The estimated transmission rates were found to be in the range of 0.01 - 4.00 per month. The result illustrated that the estimated transmission rates from the two models are different. We found that the transmission rate predicted by the model excluding the evolutionary dynamics (green line) is likely to increase after the pandemic in 2009.

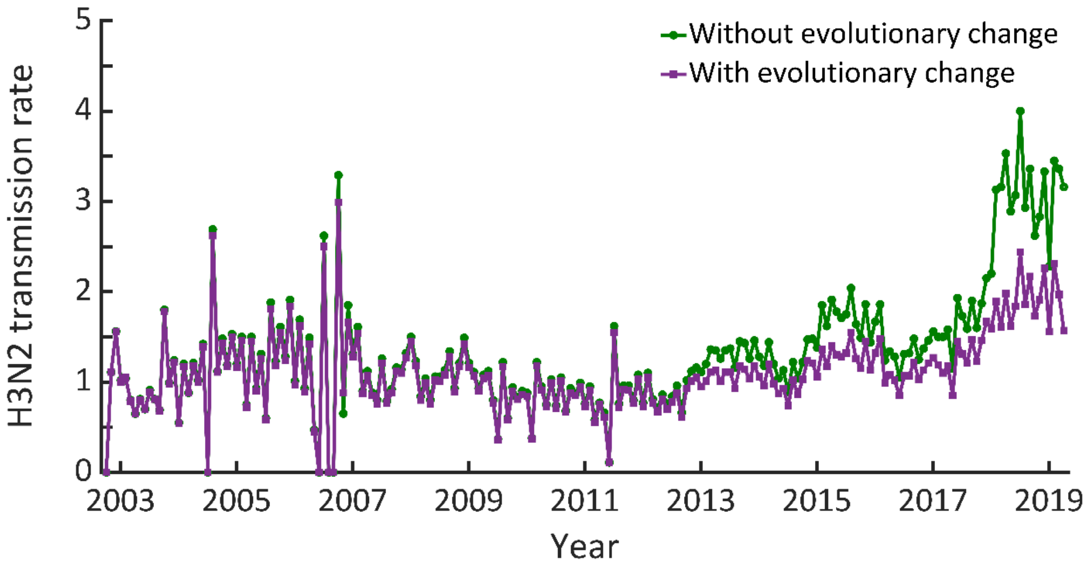


**Figure S3. Comparison of estimated monthly transmission rates.** The green line is the monthly transmission rate of the H3N2 influenza virus estimated from the model without evolutionary change. The purple line shows the monthly transmission rate of the H3N2 influenza virus estimated from the model incorporating evolutionary change of the virus.

To investigate the evolutionary competition of the A(H3N2) and A(H1N1) influenza viruses, we constructed the competitive evolutionary transmission model. **Figure S4** illustrates the fitting results of the competitive model. We found that the monthly-case trends generated by the competitive model agreed well with the reported data (R-square = 0.9998 for H3N2, and R-square = 0.9998 for H1N1). **Figure S4(C)** and **(D)** display the estimated transmission rates of the two influenza subtypes.

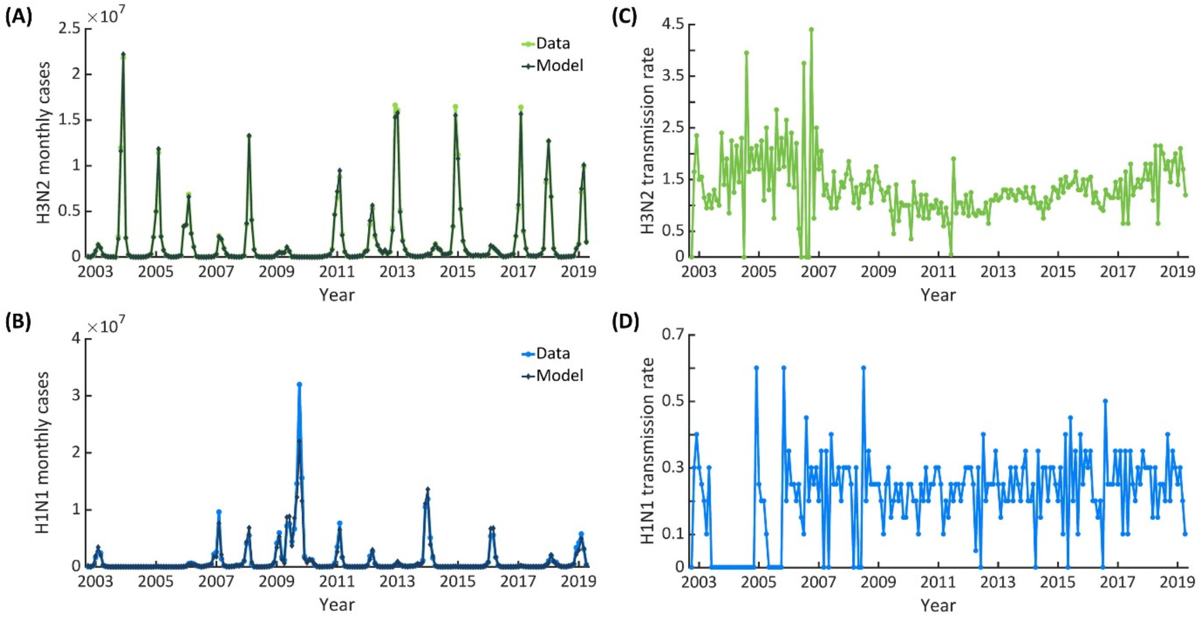


**Figure S4. The competitive model without cross-immunity. (A)** H3N2 monthly influenza cases. **(B)** H1N1 monthly influenza cases. **(C)** The time series of monthly transmission rates of influenza A(H3N2). **(D)** The time series of monthly transmission rates of influenza A(H1N1).

We also incorporated the cross-immunity between the two subtypes into the competitive model. The strength of the cross-immunity between the two strains is described by Ψ. Our model fits the observed data well with R-square = 0.9998 for A(H3N2) and R-square = 0.9998 for A(H1N1) (**Figure S5(A)** and **(B)**). The estimated monthly transmission rates which provide the best fitting of the model are shown in **Figure S5(C)** and **(D)**.

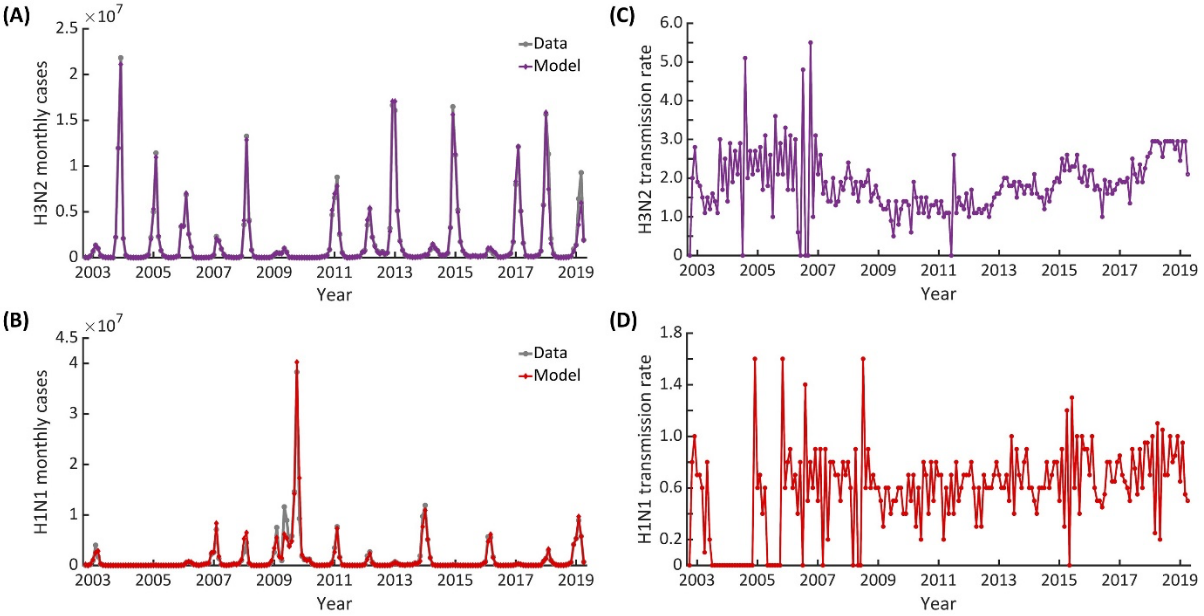


**Figure S5. The competitive model with cross-immunity. (A)** H3N2 monthly influenza cases. **(B)** H1N1 monthly influenza cases. **(C)** The time series of monthly transmission rates of influenza A(H3N2). **(D)** The time series of monthly transmission rates of influenza A(H1N1).

We used the proportion of the recovered population to measure the level of immunity in the population. The fraction of recovered individuals during epidemics of influenza A(H3N2) and A(H1N1) are shown in **Figure S6(A)** and **(B)**. Results in **Figure S6(A)** show that the differences in the level of immunity of influenza A(H3N2) tend to be stable. The immunity levels for influenza A(H3N2) predicted from three distinct models are not much different. In contrast, the proportion of the recovered population of influenza A(H1N1) in each model is remarkably different, specifically the results from the H1N1 model. We found that the single strain H1N1 model that incorporates evolutionary change predicts approximately 4-fold fewer H1N1 recovered individuals than the competitive model before and after 2009 (**Figure S6(B)**).

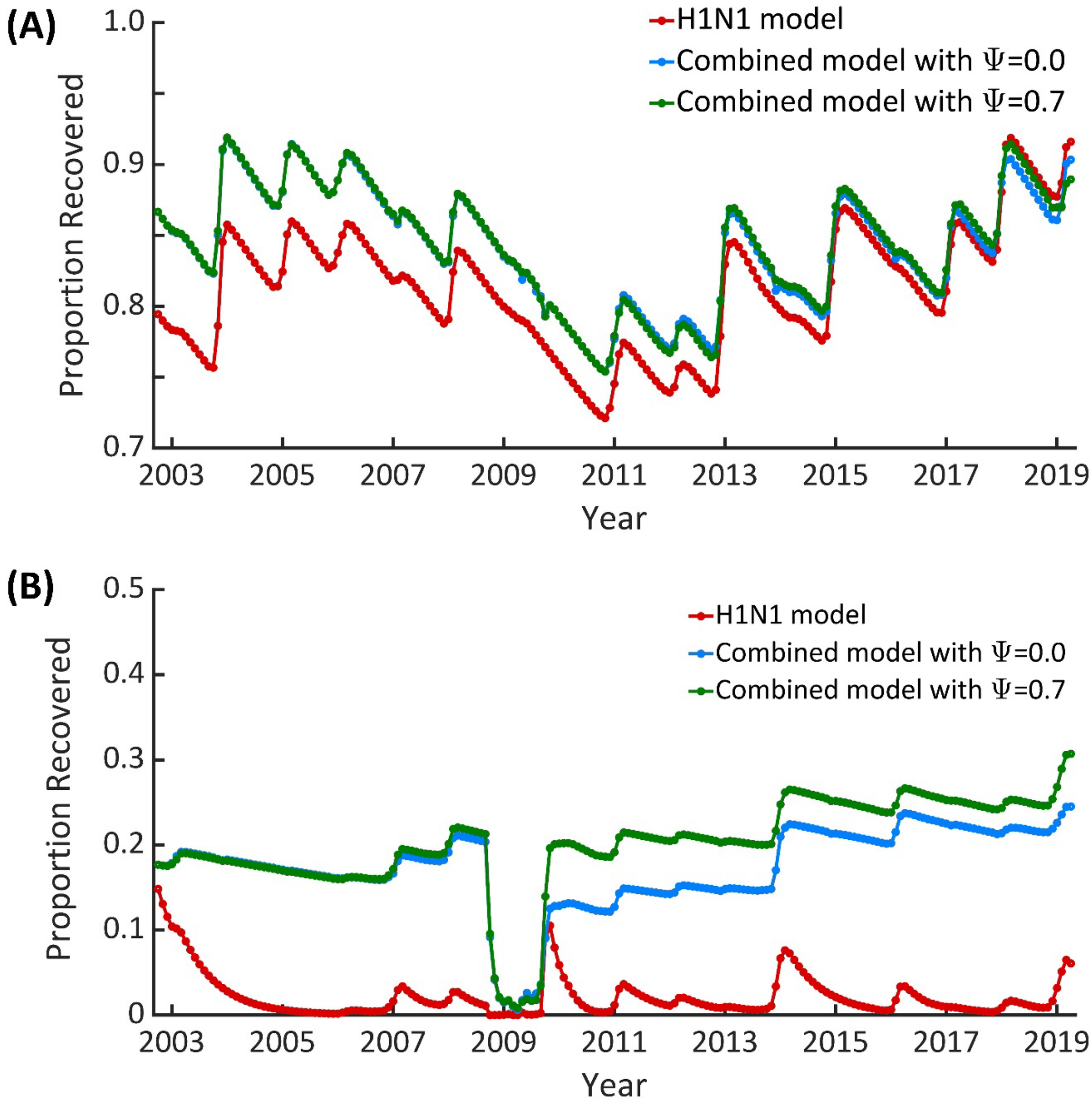


**Figure S6. The immunity level of influenza A(H3N2) and A(H1N1). (A)** The proportion of H3N2 recovered population. **(B)** The proportion of H1N1 recovered population.

**The amino acid residues of epitope regions of Influenza A(H1N1) and A(H3N2)**

For influenza A(H1N1), there are five epitope regions: Ca1, Ca2, Cb, Sa, and Sb[1-3]. The amino acid residues of epitopes of influenza A(H1N1) are summarized in **Table S1**. For influenza A(H3N2), there are five epitope regions of HA protein: A, B, C, D, and E[1,3-5]. The amino acid residues of epitopes of influenza A(H3N2) are summarized in **Table S2**.

**Table S1.** Amino acids in epitopes Sa, Sb, Ca1, Ca2, and Cb of influenza A(H1N1).

| Antigenic Site | Residue Number | Total |
|---|---|---|
| **Sa** | 124, 125, 153-157, 159-164, | 13 |
| **Sb** | 184-195 | 12 |
| **Ca1** | 166-170, 203, 204, 205, 235-237 | 11 |
| **Ca2** | 137-142, 221, 222 | 8 |
| **Cb** | 70-75 | 6 |

**Table S2.** Amino acids in epitopes A, B, C, D, and E of influenza A(H3N2).

| Antigenic Site | Residue Number | Total |
|---|---|---|
| **A** | 122, 124, 126, 130-133, 135, 137, 138, 140, 142-146, 150, 152, 168 | 19 |
| **B** | 128, 129, 155-160, 163-165, 186-190, 192-194, 196-198 | 22 |
| **C** | 44-48, 50, 51, 53, 54, 273, 275, 276, 278-280, 294, 297, 299, 300, 304, 305, 307-312 | 27 |
| **D** | 96, 102, 103, 117, 121, 167, 170-177, 179, 182, 201, 203, 207-209, 212-219, 226-230, 238, 240, 242, 244, 246-248 | 41 |
| **E** | 57, 59, 62, 63, 67, 75, 78, 80-83, 86-88, 91, 92, 94, 109, 260-262, 265 | 22 |
